# Identification and characterization of distinct cell cycle stages in cardiomyocytes using the FUCCI transgenic system

**DOI:** 10.1101/2021.08.11.455626

**Authors:** Marion Baniol, Francesca Murganti, Agata Smialowska, Joni Panula, Enikö Lazar, Viveka Brockman, Sarantis Giatrellis, Wouter Derks, Olaf Bergmann

## Abstract

Understanding the regulatory mechanism by which cardiomyocyte proliferation transitions to endoreplication and cell cycle arrest during the neonatal period is crucial for identifying proproliferative factors and developing regenerative therapies.

We used a transgenic mouse model based on the fluorescent ubiquitination-based cell cycle indicator (FUCCI) system to isolate and characterize cycling cardiomyocytes at different cell cycle stages at a single-cell resolution. Single-cell transcriptome analysis of cycling and noncycling cardiomyocytes was performed at postnatal days 0 (P0) and 7 (P7).

The FUCCI system proved to be efficient for the identification of cycling cardiomyocytes with the highest mitotic activity at birth, followed by a gradual decline in the number of cycling and mitotic cardiomyocytes during the neonatal period. Cardiomyocytes showed premature cell cycle exit at G1/S shortly after birth and delayed G1/S progression during endoreplication at P7. Single-cell RNA-seq confirmed previously described signaling pathways involved in cardiomyocyte proliferation (Erbb2 and Hippo/YAP), cardiomyocyte motility, and maturation-related transcriptional changes during postnatal development, including the metabolic switch from glycolysis to fatty acid oxidation in cardiomyocytes. Additionally, we generated transcriptional profiles specific to cell division and endoreplication in cardiomyocytes.

Deciphering transcriptional changes at different developmental stages and in a cell cycle-specific manner may facilitate the identification of genes important for adult cardiomyocyte proliferation and heart regeneration.

**Main findings:** FUCCI reliably identifies cycling cardiomyocytes at distinct cell cycle stages in neonatal, juvenile, and adult hearts.

Cell cycle activity decreases as the metabolic switch transitions from glycolysis to fatty acid metabolism in postnatal cardiomyocytes.

Cell cycle arrest at G1/S is linked to the DNA damage response in postnatal cardiomyocytes.
Distinct gene expression patterns are linked to different cell cycle phases in dividing and endoreplicating postnatal cardiomyocytes.

## Introduction

Cardiovascular diseases (CVDs) are the primary contributors to mortality and a major socioeconomic burden in the Western world (Dagenais et al., 2020). While certain pathophysiological steps might differ between disease processes covered by this umbrella term, the progressive loss of cardiomyocytes and the consequential decline in heart contractility are common features in many. Current clinical strategies target the pharmacological or mechanical support of the failing heart, while the potential replacement of the failing myocardium with new cardiomyocytes remains a challenging and unachieved goal (Chong et al., 2014).

Cardiomyocytes are renewed at low levels in the hearts of adult mammals (Ali et al., 2014; Alkass et al., 2015; Senyo et al., 2013), including humans (Bergmann et al., 2009, 2015; Mollova et al., 2013). Although the source of new cardiomyocytes was debated previously, the current consensus is that new cardiomyocytes are mainly formed from preexisting cardiomyocytes (Eschenhagen et al., 2017), providing a strong basis for the development of therapies aimed at reactivating the cell cycle in cardiomyocytes to promote de novo cell formation (Bassat et al., 2017; Liu et al., 2019a; Uva et al., 2015; Wei et al., 2015).

However, the proliferative capacity of cardiomyocytes sharply declines shortly after birth, and instead of increasing in number, they expand in size and become polyploid by increasing their DNA content in cycles of endoreplication and cytokinesis failure (Derks and Bergmann, 2020). The molecular mechanisms regulating this sharp decline in proliferation and the initiation of polyploidy and multinucleation remain largely unclear.

Detecting proliferating cardiomyocytes has been notoriously difficult and has some major technical challenges, as only a very small proportion of cardiomyocytes enter the cell cycle at any given time, and cardiomyocytes are surrounded by cell types with much higher mitotic activity (Derks and Bergmann, 2020; Hesse et al., 2018; Leone et al., 2015).

Recently, genetic cell cycle indicators proved to be powerful tools in studying the characteristics and regulation of the cardiomyocyte cell cycle (Bergmann and Braun, 2016; Jr et al., 2018; Raulf et al., 2015). We employed the fluorescence ubiquitination-based cell cycle indicator (FUCCI) system to identify and isolate cycling cardiomyocytes in postnatal and adult mice. This genetically encoded, two-color sensor allows distinct fluorescent labeling of nuclei in cycling and noncycling cells by relying on oscillating ubiquitination and subsequent degradation of the cell cycle proteins Cdt1 and geminin (Hashimoto et al., 2014, 2015; Jr et al., 2018; Sakaue-Sawano et al., 2008). This system labels nuclei of cells in the G0/G1 cell cycle phase with the red fluorescent protein monomeric Kusabira Orange (mKO2^+^) and the nuclei of cells in the S/G2/M phase with the green fluorescent protein monomeric Azami Green (mAG^+^), while cells at the G1/S transition appear double positive (mKO2^+^/mAG^+^).

We used the FUCCI system to describe the dynamics of the cell cycle activity of cardiomyocytes throughout postnatal maturation with single-cell resolution. By enriching cycling cardiomyocytes, we revealed distinct transcriptional profiles and cell cycle stage-dependent subpopulations of cells undergoing cell division/mitosis and endoreplication.

## Results and Discussion

Neonatal heart regeneration relies on cardiomyocyte proliferation following injury (Porrello et al., 2011; Xin et al., 2013), but this proliferative activity declines gradually with premature cell cycle exit, resulting in multinucleation during the first postnatal weeks. Although several postnatal changes in metabolism, immune response and neural innervation have been shown to be linked with this process (Aurora et al., 2014; Mahmoud et al., 2015; Puente et al., 2014), the exact mechanisms that regulate cell cycle exit shortly after birth remain unclear. Here, we used the cell cycle indicator FUCCI to study the cell cycle dynamics of dividing and endoreplicating single neonatal cardiomyocytes and to explore transcriptional profiles regulating cardiomyocyte cell cycle activity in the postnatal murine heart with single-cell resolution.

### FUCCI identifies cardiomyocytes in neonatal and adult mouse hearts

We first determined the expression of FUCCI in mouse heart tissue sections at postnatal days 0 (P0), P7, P15 and adulthood (P90) (Fig. 1A). To detect cardiomyocyte-specific signals, we used pericentriolar material 1 (PCM-1) as a marker for cardiomyocyte nuclei (Bergmann and Jovinge, 2012; Bergmann et al., 2011, 2015) (Fig. 1B, Suppl. Fig. 1A).

**Figure 1.**
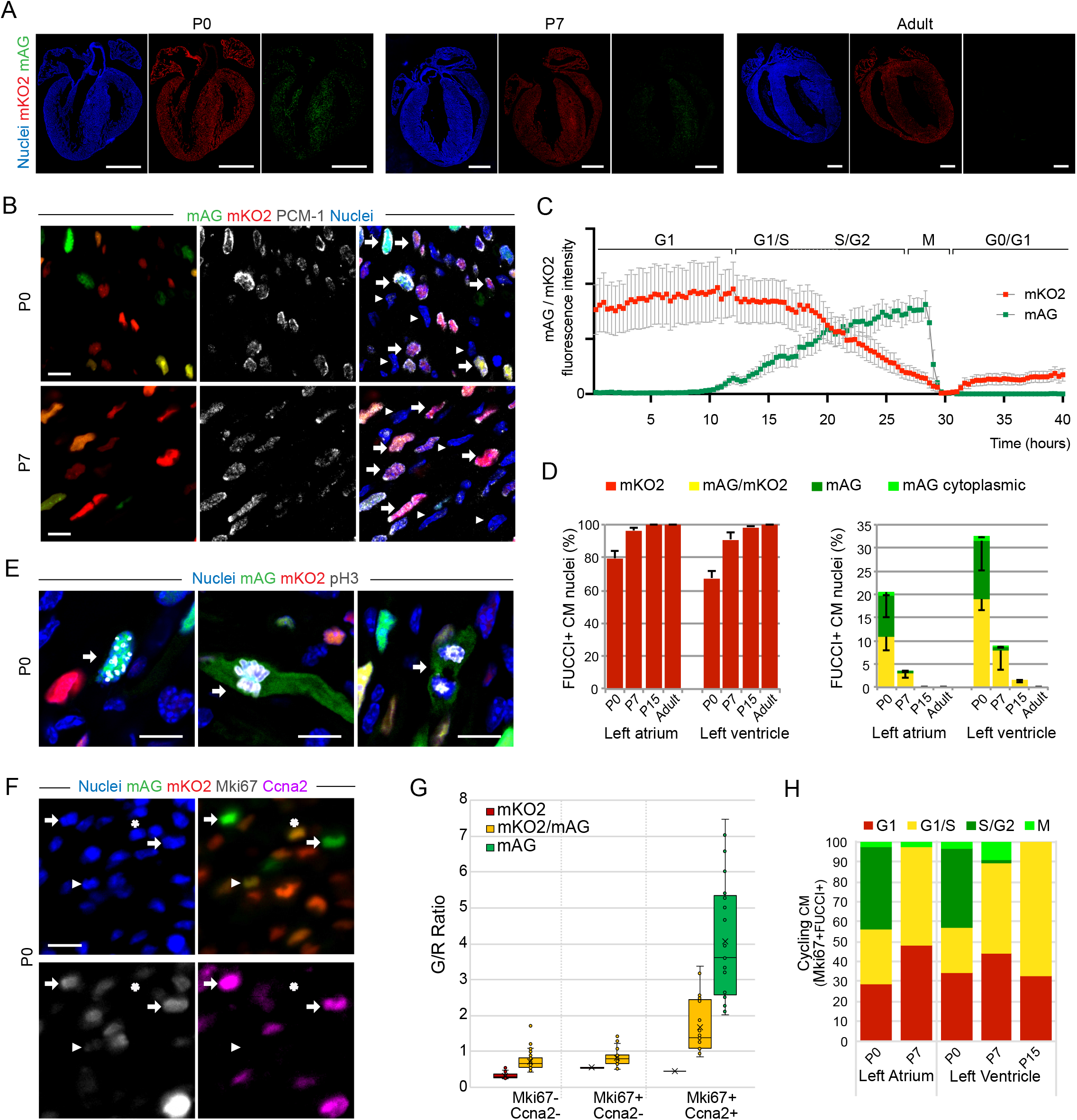
FUCCI fluorescence identifies cycling cardiomyocytes in heart tissue. (A) Longitudinal whole heart sections of FUCCI P0, P7 and adult hearts. Nuclei are stained with Hoechst. Scale bar 1 mm. (B) FUCCI expression in P0 and P7 left ventricular tissue sections used for the quantification of mKO2^+^, mAG^+^, and mKO2^+^/mAG^+^ cardiomyocyte nuclei labeled with PCM-1 (white arrows). White arrowheads indicate nonmyocyte nuclei. Nuclear FUCCI fluorescence with mKO2 and mAG is visible only in cardiomyocytes (arrows). Scale bar 10 μm. (C) Oscillation of nuclear fluorescence levels of FUCCI probes measured by live imaging of a primary culture of P0 cardiomyocytes undergoing cell division or multinucleation. The average values of 3 dividing and 7 multinucleating cardiomyocytes are shown. (D) Quantification showing the percentage of mKO2^+^ cardiomyocytes (left panel) and mKO2^+^/mAG^+^ and mAG^+^ (nuclear and cytoplasmic) cardiomyocytes (right panel) in the left atrium and left ventricle of P0, P7, P15 and adult (P90) hearts. Average values, n=3 hearts analyzed per time point, error bars represent SD. (E) Mitotic FUCCI cardiomyocytes stained with pH 3 (gray) in a P0 heart. The left picture shows cardiomyocytes in late G2 phase with nuclear localization of mAG, while the mid- and right pictures show the cytoplasmic localization of mAG after nuclear envelope breakdown in metaphase and anaphase, respectively, as confirmed by the condensed state of their DNA (Hoechst, blue). Scale bar 10 μm. (F) P0 FUCCI heart showing overlapping immunostaining of Ccna2 (magenta) and Ki67 (gray) with mAG^+^/mKO2^+^ nuclei. White arrows indicate early S-phase cardiomyocyte nuclei, white arrowhead indicates cycling G1/S cardiomyocytes, and white star indicates G1/S arrested cardiomyocytes. Scale bar 10 μm. (G) Boxplot distribution of the ratio of mAG over mKO2 mean fluorescent intensities measured in FUCCI nuclei that were found to be either noncycling (Ki67^-^Ccna2^-^), at the G1/S transition (Ki67^+^/Ccna2^-^), or in S-phase (Ki67^+^/Ccna2^+^). (H) Proportion of cycling cardiomyocytes (Ki67^+^/FUCCI^+^ nuclei) in G1, G1/S, S/G2 and M-phase in the left atrium and left ventricle of P0, P7, and P15 hearts.

Most cardiomyocytes expressed one of the FUCCI markers (exceeding 81% in P0 and 92% in P7, P15 and adult ventricular cardiomyocytes) (Suppl. Fig. 1B). Importantly, FUCCI fluorescence in the heart was the most intense in cardiomyocytes, rendering fluorescence in every other cell type almost undetectable, as previously observed in embryonic hearts (Hashimoto et al., 2014). The few FUCCI-positive nuclei detected in nonmyocytes accounted for less than 1% of the total FUCCI fluorescent nuclei observed, which were mainly present in the atria at all developmental time points and in P0 ventricular tissue. No FUCCI-positive nonmyocyte nuclei were detected out of an average of 600 nuclei analyzed per heart in the ventricles of P7, P15 and adult heart sections (Suppl. Fig. 1C). Thus, this system proved to be robust enough for the quantification of cardiomyocyte cell cycle activity and for performing high-throughput single-cell analysis of cycling cardiomyocytes in postnatal hearts.

### Isolated neonatal FUCCI cardiomyocytes undergo binucleation and cytokinesis

As previously reported, the FUCCI system can be employed for single-cell live imaging of cycling postnatal cardiomyocytes in primary cell culture (Jr et al., 2018) and in ex vivo slice culture (Hashimoto et al., 2014). We detected clear oscillation patterns of FUCCI expression in cycling cardiomyocytes over a period of 40 h (Fig. 1C) in isolated neonatal cardiomyocytes (P0). Nuclear mAG fluorescence dropped dramatically in early mitosis along with nuclear envelope breakdown in both cytokinesis and binucleation events, and the average duration of S/G2/M was 15.1 h ± 4.0 h SD for P0 cardiomyocytes undergoing cytokinesis and binucleation, with no difference in cell cycle length (unpaired t-test, p = 0.17) (Suppl. Fig. 1D upper and middle panel). Thus, FUCCI allows for the characterization of postnatal cardiomyocyte cell cycle kinetics using single-cell live imaging.

### FUCCI reliably identifies cycling cardiomyocytes at different cell cycle stages *in vivo*

To evaluate the proportion of cardiomyocytes in each phase of the cell cycle in postnatal and adult hearts, we quantified the number of cardiomyocyte nuclei displaying either mKO2^+^ (G0/G1), mKO2^+^/mAG^+^ (G1/S) or mAG^+^ (S/G2/M) fluorescence in the left atrium and the left ventricle of P0, P7, P15 and adult hearts (Fig. 1D, see Methods). Most cardiomyocytes were found to be in G0/G1, with 67.5% ± 4.1% SD mKO2^+^ nuclei in the P0 left ventricle, a fraction that increased over 90% at P7 and P15 and reached 100% in the adult ventricle (Fig. 1D, left panel). In parallel, the fraction of mAG^+^ cardiomyocytes decreased dramatically with the first postnatal week, from 12.36% (±6.35% SD) in the P0 left ventricle to less than 1% (0.83%±0.22% SD) detected at P7 (Fig. 1D, right panel). Mitotic cardiomyocytes displayed a cytoplasmic FUCCI pattern upon nuclear envelope breakdown (1% ± 0.28% SD, at P0), allowing us to distinguish cells in mitosis that entered prometaphase (Fig. 1E). All analyzed mAG^+^ cardiomyocytes were Ki67 positive (Ki67^+^) at all developmental stages analyzed (Suppl. Fig. 1E, F). In addition, 8.3% (±1% SD) of mKO2^+^ cardiomyocytes were Ki67^+^ at P0, indicating cell cycle entry (G1 phase). This number decreased to 3.5% (±1.4% SD) and 1.7% (±0.8% SD) in the P7 and P15 ventricles, respectively (Suppl. Fig. 1F), suggesting a gradual decline in cardiomyocytes entering the cell cycle.

### Postnatal cardiomyocytes withdraw from the cell cycle at the G1/S transition

We found a substantial fraction of mKO2^+^/mAG^+^ cardiomyocytes in the early neonatal period, with 19.1% (±2.5% SD) of the cardiomyocytes being observed at P0, 7.9% (±4.1% SD) at P7, and 1.48% (± 0.51% SD) in the ventricles of P15 hearts (Fig. 1D, right pannel). Most of these double-positive cardiomyocytes showed a very low mAG to mKO2 fluorescence ratio (G/R ratio) at P0, close to the ratio of mKO2^+^ cells (Suppl. Fig. 1G), indicating that they were at the early stage of the G1/S transition. Only 22.7% (±12.7% SD) and 58.8% (±29.6% SD) of them expressed Ki67 at P0 and P7, respectively, suggesting that these double-positive cardiomyocytes prematurely left the cell cycle before entering S phase (Suppl. Fig. 1F).

To further characterize the transition from G1 to S-phase, we plotted the G/R ratio of FUCCI-positive nuclei in relation to the expression of Ki67 and Ccna2, a marker of S/G2 phase (Fig. 1F, G). Most cardiomyocyte nuclei with a G/R ratio lower than 1 were either not cycling (Ki67^-^ /Ccna2^-^) or were cycling but had not yet progressed through the G1/S checkpoint (Ki67^+^/Ccna2^-^). With single-cell live imaging analysis of P0 cardiomyocytes, we identified a fraction of FUCCI cardiomyocytes that entered G1 (mAG^+^/mKO2^+^) but regressed rapidly to mAG^-^/mKO2^+^, indicating G1/S arrest without further progression to the cell cycle (Suppl. Fig. 1D, lower panel).

In addition, approximately half of the Ki67^+^ P0 cardiomyocytes were found to be in S/G2/M phase (mAG^+^), whereas the majority of the P7 and P15 Ki67^+^ cardiomyocytes were found to be at G1/S transition (mAG^+^/KO2^+^) (Fig. 1H), suggesting a delay at the transition from G1 to early S-phase during the course of the postnatal period. However, we could not find evidence for cardiomyocytes being arrested in the G1/S stage for an extended period, as reported earlier (Jr et al., 2018), because we could not detect any persistence of mAG fluorescence above the background level in cardiomyocytes after the postnatal period (Suppl. Fig. 1H).

### FUCCI allows for single cardiomyocyte transcriptomics of distinct cell cycle phases

We performed single-cell RNA sequencing of cardiomyocytes at birth (P0) and at P7 on cells sorted using the FUCCI system. Most cell cycle activity in cardiomyocytes at P0 is attributed to cell division, whereas maturing cardiomyocytes at P7 undergo mainly multinucleation (Alkass et al., 2015). Accordingly, we isolated single FUCCI cardiomyocytes from P0 and P7 whole hearts and sorted them for subsequent single-cell RNA sequencing (Fig. 2A) based on their mKO2 and mAG expression (Fig. 2B). After quality control and filtering out empty wells, we found that all but 4 of the remaining 283 cells expressed cardiomyocyte markers and were negative for other cell type markers, and the 4 outliers were retrospectively filtered out (see Methods and Suppl. Fig. 2A, B). We applied shared nearest neighbor unsupervised clustering and obtained 6 cardiomyocyte subpopulations (Fig. 2C) that were annotated based on their cell cycle status (cycling and noncycling), atrial or ventricular origin (aCM and vCM) and developmental stage (P0 and P7), as determined by the expression levels of established marker genes such as Myl2 (vCM), Myl4 (aCM) and Cdk1 (cell cycle) (Fig. 2D). Atrial CMs at P0 and P7 clustered together, suggesting limited developmental changes in their transcriptome at these stages. However, the small number of sorted P0 aCMs might have also accounted for a lack of separation. Subsequent analysis of marker genes for each cluster revealed groups of genes specific not only to atrial or ventricular CMs but also to their P0 and P7 developmental stages (Fig. 2E).

**Figure 2.**
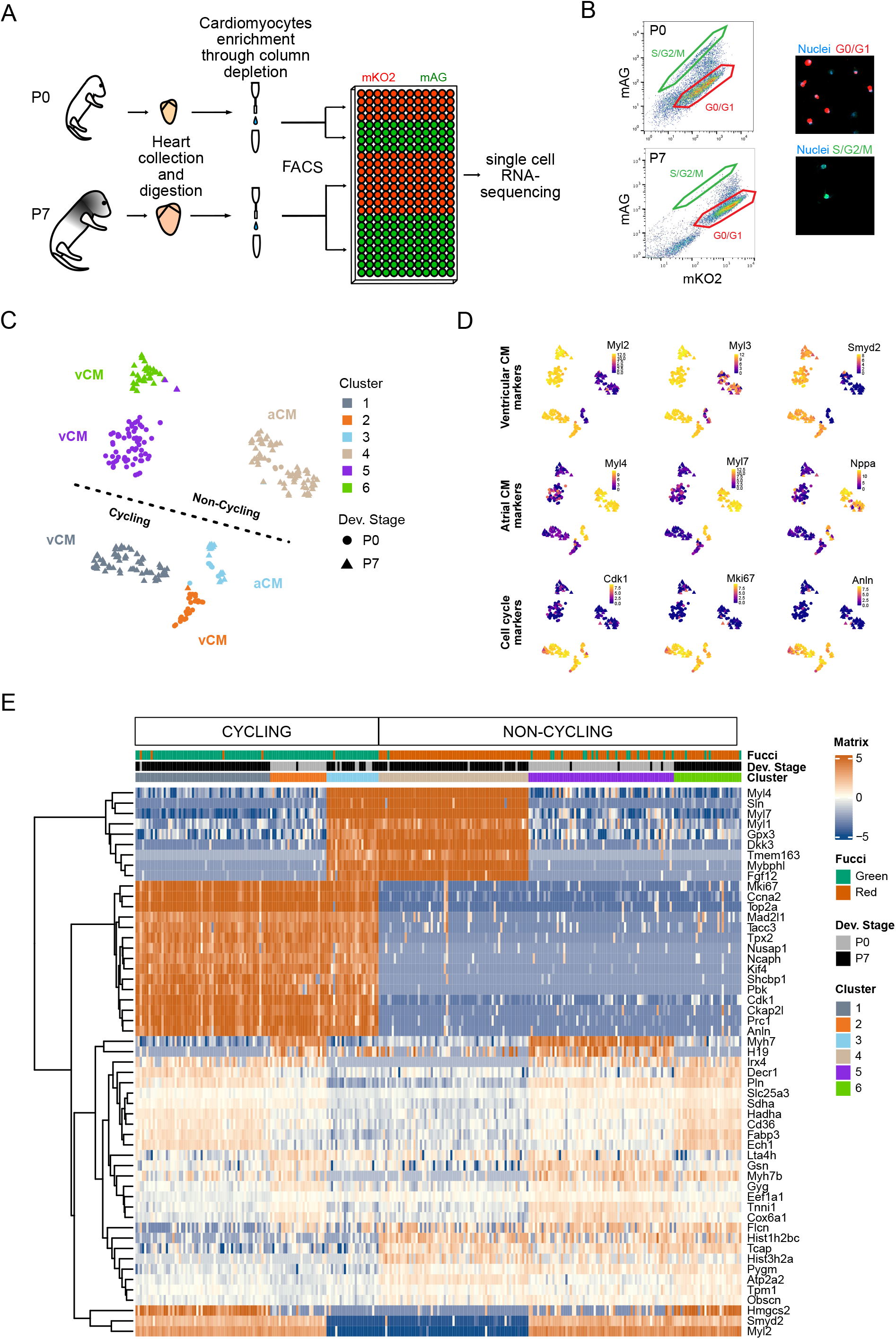
Single-cell RNA sequencing of FUCCI cardiomyocytes identifies cycling and noncycling cells of atrial and ventricular origin. (A) Schematic view of the single-cell RNA-seq experimental workflow. (B) Distribution of FUCCI fluorescence in isolated live cardiomyocytes from P0 and P7 hearts analyzed by flow cytometry. The gating strategy for the selection and downstream single-cell RNA sequencing of cycling (mAG^+^) and noncycling (mKO2^+^) cardiomyocytes is highlighted (left panel). Representative images of cells gated as mKO2^+^ and mAG^+^ (right panel). Nuclei were stained with Hoechst (blue). (C) T-SNE plot showing SNN clustering of 279 cardiomyocytes isolated from P0 and P7 FUCCI hearts. Each cell is represented by a dot shape based on its developmental stage. Cell populations were identified based on the expression of known marker genes. (D) t-SNE plots showing the expression level of known marker genes for ventricular cardiomyocytes (upper row), atrial cardiomyocytes (middle row), and the cell cycle (lower row) for each analyzed cell. Color scale represents the normalized log counts. (E) Heatmap of the expression values of the top 50 markers across cell populations. Log-normalized counts are plotted after scaling by row.

### Cardiomyocytes undergo substantial transcriptional changes during postnatal maturation

We first confirmed previously described developmental changes with differentially expressed (DE) genes between the P0-vCM and P7-vCM clusters (Quaife-Ryan et al., 2017). Analysis of P0 vs. P7 DE genes in both cycling and noncycling vCM clusters showed a common pattern of P7 upregulated genes associated with GO terms of lipid metabolism (Hmgcs2, Fabp3, Pdk4), ATP production (Ucp2) and heart contraction (Fig. 3A-C), all indicative of increased cardiomyocyte maturation (Karbassi et al., 2020). We also found genes that have been described to be involved in cell cycle arrest (p38, Ccng1). In particular, cyclin G1 (Ccng1) was previously shown to play a role in G2/M arrest, damage recovery and growth promotion after cellular stress (Kimura et al., 2001). More importantly, it was previously linked to the development of polyploidy in cardiomyocytes (Liu et al., 2010). DE genes in P0-vCMs, on the other hand, were associated with GO terms of glucose metabolism, cell growth and cardiomyocyte proliferation (Igf2, H19, Fstl1, Tbx20), indicative of a relatively immature state (Karbassi et al., 2020). Additionally, the high representation of cell motility and cell migration-related terms highlight the regenerative capacity of P0 hearts, as epithelial-to-mesenchymal transition (EMT) of cardiomyocytes and migration to injury sites have been shown to be involved in heart repair (Aharonov et al., 2020). In this context, we observed a switch in integrin alpha (Itga) subunits, with Itga1 being upregulated in P0-vCMs, while Itga7 was upregulated in P7-vCMs. The Itga1 subunit, which is a cell surface receptor for laminin and collagen, has been linked to cell migration in embryonic fibroblasts, suggesting a role in cardiomyocyte migration to the injury site in the neonate regenerative heart (Gardner et al., 1996).

**Figure 3.**
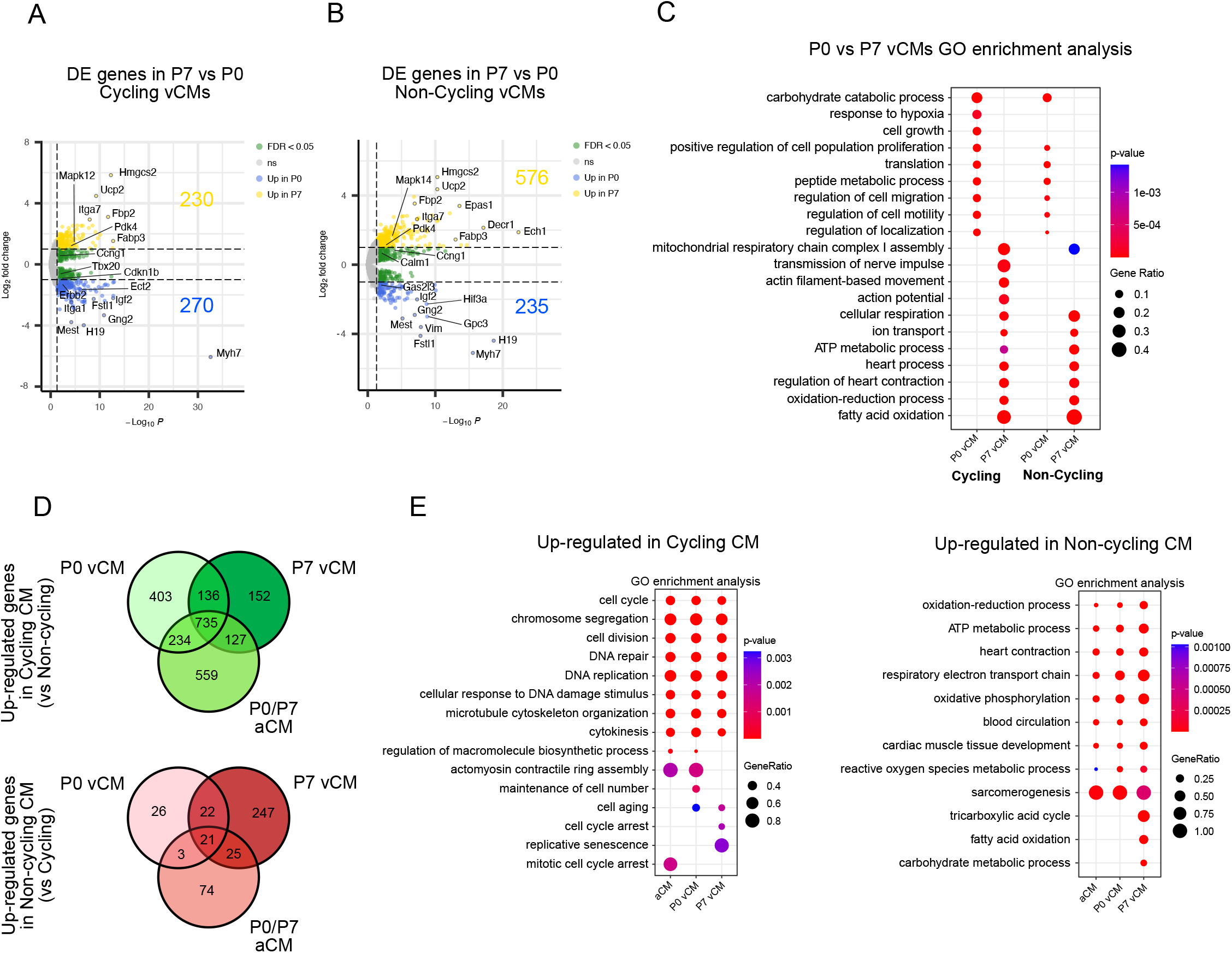
Distinct gene expression profiles of P0 and P7 ventricular cardiomyocytes. (A) Volcano plot showing the number of significantly upregulated (yellow) and downregulated (blue) genes in cycling vCMs at P7 compared to P0. (B) Volcano plot showing the number of significantly upregulated (yellow) and downregulated (blue) genes in noncycling vCMs at P7 compared to P0. (C) Enriched GO terms for genes differentially expressed between P0 and P7 vCMs for cycling and noncycling vCM clusters. (D) Venn diagrams showing the number of shared and uniquely differentially expressed genes in the cycling P0-vCM, P7-vCM and P0/P7-aCM clusters (upper panel) compared to their noncycling counterparts (lower panel). (E) Enriched GO terms for genes upregulated in cycling cardiomyocytes (left panel) and in noncycling cardiomyocytes (right panel) of all three subpopulations.

Upregulated genes in cycling P0-vCMs included members of the Hippo/YAP signaling pathway and Nrg1/ErbB2 signaling pathway (Erbb2, Hey2, Amotl1) and genes known to be required for cardiomyocyte cytokinesis (e.g., Ect2 and RhoA)(González-Rosa et al., 2018; Liu et al., 2019b; Stopp et al., 2017).

To interrogate what distinguishes cycling from noncycling cardiomyocytes, we performed differential gene expression analysis between cycling and noncycling clusters for all three cardiomyocyte subpopulations (P0-vCM, P7-vCM and P0/P7-aCM) (Fig. 3D and E). Cell cycle exit involved the downregulation of over a thousand genes (1508, 1150 and 1655 genes with a fold change >2 in P0-vCM, P7-vCM, and P0/P7-aCM, respectively), with large overlap between all three populations (Fig. 3D), suggesting a core cell cycle transcriptional program. Genes upregulated in noncycling cells were less abundant and more variable between the cardiomyocyte subpopulations. GO analysis of these DE genes showed that processes enriched in cycling populations involved the cell cycle, cell division and DNA replication as well as DNA repair and DNA damage responses, whereas noncycling CMs expressed genes important for energy production, mitochondrial activity, and heart contraction, consistent with the functional role of cardiomyocytes (Fig. 3E).

### Reclustering of cycling cardiomyocytes identifies distinct cell cycle phases in postnatal cardiomyocytes

The heterogeneous distribution of cell cycle gene expression levels across the entire cycling cardiomyocyte population suggested that these cells were in distinct stages of the cell cycle (Fig. 2D, Suppl. Fig. 2C). We therefore subjected the cycling cardiomyocytes to hierarchical clustering (Fig. 4A) and plotted the expression levels of reported marker genes for G1/S transition, S-phase, and G2/M-phase to assign the obtained clusters to these cell cycle phases (Fig. 4B).

**Figure 4.**
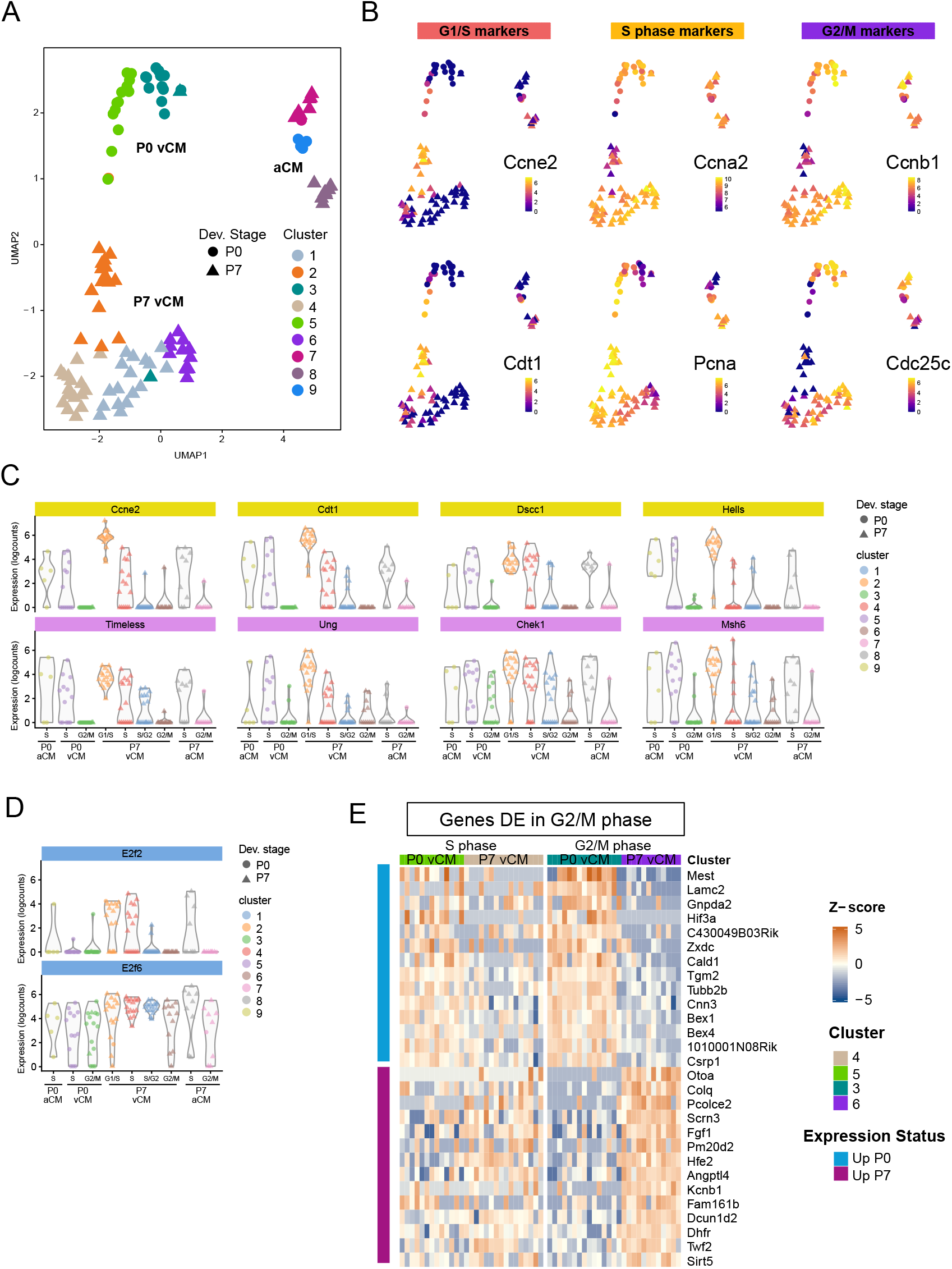
Analysis of distinct cell cycle phases in neonatal cardiomyocytes. (A) UMAP plot of cycling cardiomyocytes colored by cluster. (B) UMAP plots showing the expression level of representative marker genes of G1/S, S and G2/M phase. Color scale represents the normalized log counts. (C) Violin plot showing the expression levels of genes involved in the G1/S transition (yellow) and in the DNA damage response (pink) for each cluster. (D) Violin plot showing E2F transcription factor expression levels for each cluster. (E) Heatmap plot of the top differentially expressed genes between P0 and P7-vCM clusters specifically at G2/M phase (right panel).

While most clusters were related to either S-phase or G2/M cardiomyocytes, Cluster 2 (Fig. 4A, orange) identified a distinct P7-vCM population characterized by the expression of genes such as Cyclin E2 and Cdt1, corresponding to the G1/S transition (Barr et al., 2016; Pozo and Cook, 2016). Although these genes are required for normal cell cycle activity in all developmental stages, we found most Cyclin E2-expressing cardiomyocytes at P7 within Cluster 2 (Fig. 4B). Most cycling P7 cardiomyocytes were in G1/S and S-phase (75%) compared to cycling P0 cardiomyocytes (50%), where we found that an equal number had already progressed to G2/M (Fig. 1). A delay of G1/S transition and S-phase progression is further supported by our imaging analysis of neonatal FUCCI hearts, where we found the largest fraction of mAG^+^/mKO^+^ at P7 (Fig. 1). An increased length of the S/G2/M phase duration in cardiomyocytes has been previously observed from the embryonic to neonatal stage (Hashimoto et al., 2014),thus appearing to continue during postnatal development.

In addition to G1/S transition genes, we found several DNA damage response (DDR) and DNA damage checkpoint genes, such as Timeless, Ung and Chek1, to be upregulated in Cluster 2 (P7-vCMs in G1/S) (Fig. 4C), consistent with DDR reported to be caused by ROS production with an increase in oxidative stress (Cardoso et al., 2020; Puente et al., 2014)

Many of the genes upregulated in Cluster 2 were previously described as upregulated in P7 mononucleated cardiomyocytes compared to binucleated cardiomyocytes and as regulated by the E2F transcription factor family (Windmueller et al., 2020).

Thus, we analyzed the expression levels of E2f genes in all our cardiomyocyte populations. We found that E2f1, E2f2, E2f7, and E2f8 were expressed only in cycling cardiomyocytes (Suppl. Fig. 3B), with E2f1 being mostly expressed at G1/S and early S-phase and E2f7 and Ef8 being expressed throughout the cell cycle (Suppl. Fig. 3C), consistent with previous reports (Cuitiño et al., 2019; Ebelt et al., 2005, 2006)

E2f2 expression was shown in proliferating cells throughout the cell cycle (Cuitiño et al., 2019); however, we found that its expression was restricted almost exclusively to cycling P7 vCMs, particularly to cardiomyocytes in Cluster 2 (G1/S) (Fig. 4D), suggesting that E2f2 acts as a positive regulator in cell cycle progression in mononucleated cycling cardiomyocytes undergoing endoreplication.

E2f6 was found to be expressed in all P7 cardiomyocytes independent of the cell cycle state, including noncycling cells (Fig. 4D, Suppl. Fig. 3B). As a repressor of other E2f genes, particularly E2f1, and their targets, E2f6 is thought to act both as an inhibitor of cell cycle reentry and as an anti-apoptotic factor (Cartwright et al., 1998; Gaubatz et al., 1998).

Next, we analyzed the expression profiles of G2/M-annotated P0 and P7 cardiomyocytes, revealing a subset of genes differentially expressed between P0 and P7 G2/M cardiomyocytes (Fig. 4E, Suppl. Fig. 3D). We observed a switch of tubulin isoforms from Tubb2b to Tubb6 and Tuba4a in neonatal G2/M cardiomyocytes (Suppl. Fig. 3D, E). Tuba4a is expressed mostly in the adult heart, while Tubb2b is most abundantly expressed in embryonic tissues (Yue et al., 2014). While several tubulins are involved in the mitotic cell cycle, the exact physiological relevance of this isoform switch and its potential impact on cytokinesis remain to be determined. Additionally, along with Ect2, which is a known regulator of cytokinesis (González-Rosa et al., 2018; Liu et al., 2019b), several genes related to cell motility and the actomyosin cytoskeleton were downregulated in P7-vCMs, such as Lamc2, Cald1, Cnn3, Bex4, and Csrp1, providing potential new leads on factors involved in successful cytokinesis in cardiomyocytes. Of note, the imprinted gene Mest, which is globally upregulated in P0-vCMs, showed its highest differential expression in cycling G2/M cardiomyocytes (Suppl. Fig. 3E). Although its function remains largely unknown, Mest has been found to be expressed extensively in fetal tissues, and its loss of imprinting and overexpression was found to be associated with cancer (Moon et al., 2010; Pedersen et al., 1999). Mest knockout mice showed trabeculation disturbances, with an increased thickness and reduced density of the compact myocardium in the embryonic heart, a feature resembling human ventricular noncompaction cardiomyopathy (King et al., 2002). However, the precise role of Mest in the regulation of cardiomyocyte proliferation and heart regeneration still needs to be determined in future studies.

In summary, the FUCCI system utilized in the present investigation is a valuable tool for studying cell cycle activity decreases and gene expression changes in neonatal cardiomyocytes at distinct cell cycle phases at the single-cell level. In particular, these results support previous findings, such as the activation of signaling pathways (Erbb2 and Hippo/YAP) in neonatal cardiomyocytes and metabolic changes associated with the maturation process of cardiomyocytes. In addition, we found changes in gene expression linked to cell motility and cytoskeleton composition that might play a role in cytokinesis failure. Moreover, we found evidence for cardiomyocyte cell cycle arrest at G1/S, which was linked to the DNA damage response in postnatal cardiomyocytes. Furthermore, our results suggest that E2f2 transcription factor expression might be crucial for cell cycle progression in endoreplicating postnatal cardiomyocytes.

## Methods

### Animals

Animals were housed in the Wallenberg Laboratory animal facility on a 12-hour light/dark cycle and were provided food and water ad libitum. All breeding and organ collection protocols were performed in accordance with the Swedish and European Union guidelines and approved by the institutional ethical committee (Stockholms Norra Djurförsöksetiska Nämnd). FUCCI transgenic mice (#504: Fucci-S/G2/M-Green (mAG-hGeminin (1/110)); #596: Fucci-G1-Red (mKO2-hCdt1(30/120)) (Sakaue-Sawano et al., 2008) were obtained from the RIKEN BioResource Research Center. For heart collection, neonatal mice (P0, P7) were euthanized by decapitation, whereas adolescent (P15) and adult mice (P90) were euthanized using CO2.

### Immunohistochemistry

Hearts were collected in PBS, fixed for 2 h in 4% PFA at 4 °C, cryoprotected in 30% sucrose overnight, embedded in OCT compound and frozen on dry ice (P0-P15) or in isopentane cooled with dry ice (adult). Cryosections were washed in PBS and incubated in blocking buffer (4% NDS (normal donkey serum) and 0.1% Triton X-100 in PBS) overnight at room temperature with the following primary antibodies: anti-PCM1 (1:200, rabbit, Atlas HPA023374), anti-Ki67 (1:100, rat, eBioscience 14-5698-82), anti-phospho-histone 3 (1:200, rabbit, Santa Cruz), antige-minin (1:1000, rabbit, Abcam ab175799), anti-alpha-actinin (1:200, mouse, Sigma A7811), and anti-Ccna2 (1:200, rabbit, Atlas HPA020626). After several washes, fluorescent labeling was performed using secondary antibodies (AF647- and Cy3-conjugated anti-rabbit IgGs, Cy5-conjugated anti-rat IgG, AF647-anti-mouse IgG, (1:500, Jackson Immunoresearch Laboratories), and AF750-anti-rabbit IgG (1:500, Invitrogen A21039) in PBS for 1 h at room temperature. Nuclei were stained with Hoechst 33342 (1:2000, Molecular Probes H3570) for 30 min, washed and mounted using ProLong Gold Mounting Medium (Invitrogen). Imaging of

PCM-1, Ki67, pH3, and Geminin immunostaining of FUCCI hearts was performed using a Zeiss LSM880 confocal microscope. A Zeiss AxioScan Z1 Slide Scanner was used to image the FUCCI channels along with Hoechst 33421/Ki67/Ccna2 (5 channels). Image analysis and quantifications were performed using ImageJ-Fiji software. Individual cardiomyocyte nuclei were grouped according to their FUCCI fluorescence (mAG+; mKO2+; mAG+/mKO2+) manually. Fluorescence intensities were measured from individual nuclei from both the red and green channel with ImageJ-Fiji software and the green/red (G/R) ratios were determined accordingly.

### Primary cardiomyocyte culture and live imaging

P0 cardiomyocytes were isolated using the Mouse Neonatal Cardiomyocytes Dissociation kit (Miltenyi) following the manufacturer’s instructions. For time-lapse imaging, cells were seeded in plating medium (20% M199 (Medium 199), 65% DMEM (Dulbecco’s modified Eagle’s medium), 5% FBS (fetal bovine serum), 10% HS (horse serum)) at a seeding density of 0.15 million cells in a 24-well plate (Cellvis) coated with fibronectin (Sigma–Aldrich). Cells were allowed to recover for 1 day in an incubator (37 °C, 5% CO2). Before imaging, the cells were washed 3X with PBS, and plating medium was applied. Cell cycle progression was observed every 20 min for 72 h using a Keyence BZ X800 fluorescence microscope (Keyence, Japan). Keyence image measurement and analysis software (Keyence, Japan) and ImageJ-Fiji software were used to quantify the mAG and mKO2 fluorescent intensities. Briefly, the nucleus was segmented using the brightfield signal, and a region around the cell was defined as the background region. The intensities of mAG and mKO2 were detected in both the nucleus and background region, and the background signal was subtracted from the nuclear intensity to reduce the variability of the signal. Finally, mAG and mKO2 fluorescent intensities were plotted using GraphPad Prism over a period of 40 h. For each graph, single cells were aligned based on the drop in the mAG intensity.

### Cardiomyocyte isolation and FACS

P0 and P7 cardiomyocytes were isolated using a Mouse Neonatal Cardiomyocytes Dissociation kit (Miltenyi) following the manufacturer’s instructions. Dead cells were labeled with SYTOX Red (Invitrogen).

For flow cytometry analysis, hearts from double-transgenic FUCCI pups were selected; atria and ventricles were dissected and digested separately. For single-cell RNA sequencing, P0 cardiomyocytes were isolated from whole hearts of 11 pups from the same litter (including animals carrying only one of the transgenes), whereas P7 cardiomyocytes were isolated from 6 hearts of double-transgenic FUCCI pups only. As a result, although the FUCCI fluorescence of the sorted cardiomyocytes corresponded to their cell cycle status at the transcriptional level overall, a small but visible fraction of P0 mAG^+^ cardiomyocytes clustered as noncycling cells, which was most likely explained by the lack of the FUCCI Red transgene.

Cardiomyocyte analysis and sorting were performed on a BD Bioscience Influx flow cytometer using a 100 μm nozzle.

### Library preparation and single-cell RNA-seq

The sequencing libraries were prepared in 384-well plates using the Smart-seq2 protocol(Picelli et al., 2013), with some minor adaptations. Sequencing was performed on a HiSeq2500 instrument with a 50 bp single read setup.

### Single-cell RNA-seq data analysis

The samples were analyzed by first demultiplexing the fastq files using deindexer (https://github.com/ws6/deindexer) using nextera index adapters and a 384-well layout. Individual fastq files were then mapped to the relevant genome assembly using STARaligner (Dobin et al., 2013) using 2-pass alignment to improve the performance of de novo splice junction reads and filtered for only uniquely mapping reads that were saved in the BAM file. The expression values were computed as reads per kilobase of gene model and million mappable reads (RPKMs) to normalize for varying sequencing depths across sequenced cells and the gene lengths. The expression values were computed per gene as described in Ramsköld et al. (Ramsköld et al., 2009) using uniquely aligned reads and correcting for the uniquely alignable positions using MULTo (Storvall et al., 2013). The count matrix obtained shows the individual counts aligned to each gene per cell.

### Cell filtering and normalization

Low-quality cells were removed based on the cutoff of 3 median absolute deviations (MADs) from the median for the dataset for the following metrics: (i) library size lower than the cutoff; (ii) number of reads mapped to ERCC higher than the cutoff; (iii) number of counts to total features lower than the cutoff; and (iv) number of reads mapped to mitochondrially encoded genes higher than the cutoff. Data were normalized using the deconvolution method (Lun et al., 2016). Following normalization, features with low expression were filtered from the dataset. The relationships between cells were visualized by constructing t-stochastic neighbor embedding (t-SNE) plots for the first two components.

### Cell clustering

Cell clustering was achieved using the shared nearest neighbor (SNN) method on the whole dataset and hierarchical clustering on the cycling subset.

### Differential gene expression analysis and Gene Ontology

Marker genes for subpopulations were identified using pairwise t-tests and Wilcoxon rank-sum tests for each cluster pair as implemented in function findMarkers from the R package scran (Lun et al., 2016). In all cases, the test was to identify genes upregulated and downregulated in each pairwise comparison. For each gene and cluster, the summary effect size was defined as the effect size from the pairwise comparison with the lowest p value. The combined p value was computed by applying Simes’ method to all p values.

GoSeq (Young et al., 2010) was used to compute gene length bias in the detection of differential expression and its effect on the identification of overrepresented functional categories. The gene-to-functional category bindings were retrieved from the most recent versions of organism annotation packages available from Bioconductor.

## Acknowledgments

O.B. was supported by the Center for Regenerative Therapies Dresden, the Karolinska Institute, the Swedish Research Council, the Ragnar Söderberg Foundation, the Åke Wiberg Foundation, and the LeDucq Foundation. The single-cell transcriptome data were generated at the Eukaryotic Single-cell Genomics facility at Science for Life Laboratory in Stockholm, Sweden. We thank Karolinska Biomedicum Imaging Core, and in particular Florian Salomons for his help with image acquisition and analysis.

## Author Contributions

Conceptualization, M.B. and O.B.; Methodology, M.B., F.M., J.P., V.B., S.G.; Software, M.B. and A.S.; Investigation, M.B., F.M., J.P., S.G., E.L., W.D., and O.B.; Writing – Original Draft, M.B. and O.B.; Writing – Review & Editing, M.B., E.L., W.D. and O.B.; Supervision, W.D., M.B. and O.B.; Funding Acquisition, O.B.

---

**Supplementary Figure 1**

(A) Split images of representative immunostaining of FUCCI expression in cardiomyocytes of P0 and P7 hearts (see Fig. 1C), with additional pictures of P15 and adult left ventricular tissue sections. Cardiomyocyte nuclei are labeled with PCM-1 (gray), white arrowheads indicate mAG^+^ nuclei, and white arrows indicate double-positive nuclei. Scale bar 10 μm. (B) Quantification of the proportion of cardiomyocyte nuclei (PCM-1^+^) that were either FUCCI positive (FUCCI^+^) or FUCCI negative (FUCCI^-^) in left atrial and left ventricular tissue sections of P0, P7, P15 and adult hearts. (C) Quantification of the proportion of FUCCI^+^ nuclei that were either cardiomyocytes (PCM-1^+^ nuclei, CM) or nonmyocytes (PCM-1^-^ nuclei, non-CM). (D) Oscillation of nuclear fluorescence levels of FUCCI probes in a primary culture of P0 cardiomyocytes undergoing cell division (upper panel, average values of n=3), multinucleation (middle panel, average values of n=7), or regression to G0 at the G1/S transition (lower panel, average values of n=6). (E) Representative images of Ki67 immunostaining (gray) performed in P0, P7, and P15 FUCCI heart sections to analyze the cell cycle status of mKO2^+^, mAG^+^ and mKO2^+^/mAG^+^ nuclei. White arrowheads indicate mAG^+^ nuclei, white arrows indicate mKO2^+^/mAG^+^ nuclei, and pink arrowheads indicate mKO2^+^ nuclei. Scale bar 10 μm. (F) Quantification of the percentage of mKO2^+^, mAG^+^ and mKO2^+^/mAG^+^ nuclei that were found to be Ki67^+^ in the left atrium and left ventricle of P0, P7 and P15 hearts. All quantification data are represented as the mean ± SD, with n=3 hearts analyzed for each time point. (G) Distribution of the ratio of mAG over mKO2 mean fluorescent intensities measured in P0 FUCCI nuclei. (H) Immunostaining of the expression of endogenous Geminin (magenta) in P0, P7, P15 and adult FUCCI hearts expressing only the mAG-hGeminin(1/110) transgenic construct. Scale bar 20 μm.

**Supplementary Figure 2**

(A) Quality control of the single-cell RNA-seq data after filtering out empty cells. Control features included the distribution of library size (in millions of reads detected, left panel), number of expressed genes per cell (center-left panel), proportion of ERCC spikes (center-right panel), and proportion of mitochondrial genes detected (right panel). (B) Heatmap of the normalized log counts for cardiomyocyte, endothelial cell, and fibroblast marker genes obtained from an uncensored single-cell dataset. (C) t-SNE visualization of cardiomyocytes colored by the expression levels of endogenous Cdt1 and geminin, Pcna, and Cdc25c. Color scale represents the normalized log counts.

**Supplementary Figure 3**

(A) UMAP plot of cycling cardiomyocytes colored by the expression levels of ventricular and atrial cardiomyocyte markers. Color scale represents the normalized log counts. (B) Violin plots showing the expression levels of E2F transcription factors in cycling and noncycling P0-vCMs and P7-vCMs. (C) Violin plots showing the expression levels of E2F transcription factors for each cluster of cycling cardiomyocytes. (D) Heatmap plot of the top differentially expressed genes between P0 and P7-vCM clusters in both S- and G2/M phase (left panel) and in S-phase only (right panel). (E) UMAP plot of cycling cardiomyocytes colored by the expression levels of genes differentially expressed in G2/M phase. Color scale represents the normalized log counts.

## References

Aharonov, A., Shakked, A., Umansky, K.B., Savidor, A., Genzelinakh, A., Kain, D., Lendengolts, D., Revach, O.Y., Morikawa, Y., Dong, J., et al. (2020). ERBB2 drives YAP activation and EMT-like processes during cardiac regeneration. Nat. Cell Biol. 22, 1346–1356.

Ali, S.R., Hippenmeyer, S., Saadat, L. V, Luo, L., Weissman, I.L., and Ardehali, R. (2014). Existing cardiomyocytes generate cardiomyocytes at a low rate after birth in mice. Proc. Natl. Acad. Sci. U. S. A. 111, 8850–8855.

Alkass, K., Panula, J., Westman, M., Wu, T.-D., Guerquin-Kern, J.-L., and Bergmann, O. (2015). No Evidence for Cardiomyocyte Number Expansion in Preadolescent Mice. Cell 163, 1026–1036.

Aurora, A.B., Porrello, E.R., Tan, W., Mahmoud, A.I., Hill, J.A., Bassel-Duby, R., Sadek, H.A., and Olson, E.N. (2014). Macrophages are required for neonatal heart regeneration. J Clin Invest 124, 1382–1392.

Barr, A.R., Heldt, F.S., Zhang, T., Bakal, C., and Novák, B. (2016). A Dynamical Framework for the All-or-None G1/S Transition. Cell Syst. 2, 27–37.

Bassat, E., Mutlak, Y.E., Genzelinakh, A., Shadrin, I.Y., Umansky, K.B., Yifa, O., Kain, D., Rajchman, D., Leach, J., Bassat, D.R., et al. (2017). The extracellular matrix protein agrin promotes heart regeneration in mice. Nature 547, 179–184.

Bergmann, O., and Braun, T. (2016). Caught Red-Handed. Circ. Res. 118, 3–5.

Bergmann, O., and Jovinge, S. (2012). Isolation of cardiomyocyte nuclei from post-mortem tissue. J. Vis. Exp.

Bergmann, O., Bhardwaj, R.D., Bernard, S., Zdunek, S., Barnabé-Heider, F., Walsh, S., Zupicich, J., Alkass, K., Buchholz, B.A., Druid, H., et al. (2009). Evidence for cardiomyocyte renewal in humans. Science 324, 98–102.

Bergmann, O., Zdunek, S., Alkass, K., Druid, H., Bernard, S., and Frisén, J. (2011). Identification of cardiomyocyte nuclei and assessment of ploidy for the analysis of cell turnover. Exp. Cell Res. 317, 188–194.

Bergmann, O., Zdunek, S., Felker, A., Salehpour, M., Alkass, K., Bernard, S., Sjostrom, S.L., Szewczykowska, M., Jackowska, T., dos Remedios, C., et al. (2015). Dynamics of Cell Generation and Turnover in the Human Heart. Cell 1566–1575.

Cardoso, A.C., Lam, N.T., Savla, J.J., Nakada, Y., Pereira, A.H.M., Elnwasany, A., Menendez-Montes, I., Ensley, E.L., Bezan Petric, U., Sharma, G., et al. (2020). Mitochondrial substrate utilization regulates cardiomyocyte cell-cycle progression. Nat. Metab. 2, 167–178.

Cartwright, P., Müller, H., Wagener, C., Holm, K., and Helin, K. (1998). E2F-6: A novel member of the E2F family is an inhibitor of E2F-dependent transcription. Oncogene 17, 611–623.

Chong, J.J.H., Yang, X., Don, C.W., Minami, E., Liu, Y.-W., Weyers, J.J., Mahoney, W.M., Van Biber, B., Palpant, N.J., Gantz, J. a, et al. (2014). Human embryonic-stem-cell-derived cardiomyocytes regenerate non-human primate hearts. Nature.

Cuitiño, M.C., Pécot, T., Sun, D., Kladney, R., Okano-Uchida, T., Shinde, N., Saeed, R., Perez-Castro, A.J., Webb, A., Liu, T., et al. (2019). Two Distinct E2F Transcriptional Modules Drive Cell Cycles and Differentiation. Cell Rep. 27, 3547–3560.e5.

Dagenais, G.R., Leong, D.P., Rangarajan, S., Lanas, F., Lopez-Jaramillo, P., Gupta, R., Diaz, R., Avezum, A., Oliveira, G.B.F., Wielgosz, A., et al. (2020). Variations in common diseases, hospital admissions, and deaths in middle-aged adults in 21 countries from five continents (PURE): a prospective cohort study. Lancet 395, 785–794.

Derks, W., and Bergmann, O. (2020). Polyploidy in Cardiomyocytes: Roadblock to Heart Regeneration? Circ Res 126, 552–565.

Dobin, A., Davis, C.A., Schlesinger, F., Drenkow, J., Zaleski, C., Jha, S., Batut, P., Chaisson, M., and Gingeras, T.R. (2013). STAR: Ultrafast universal RNA-seq aligner. Bioinformatics 29, 15–21.

Ebelt, H., Hufnagel, N., Neuhaus, P., Neuhaus, H., Gajawada, P., Simm, A., Müller-Werdan, U., Werdan, K., and Braun, T. (2005). Divergent siblings: E2F2 and E2F4 but not E2F1 and E2F3 induce DNA synthesis in cardiomyocytes without activation of apoptosis. Circ. Res. 96, 509–517.

Ebelt, H., Liu, Z., Müller-Werdan, U., Werdan, K., and Braun, T. (2006). Making omelets without breaking eggs: E2F-mediated induction of cardiomyoycte cell proliferation without stimulation of apoptosis. Cell Cycle 5, 2436–2439.

Eschenhagen, T., Bolli, R., Braun, T., Field, L.J., Fleischmann, B.K., Frisen, J., Giacca, M., Hare, J.M., Houser, S., Lee, R.T., et al. (2017). Cardiomyocyte Regeneration: A Consensus Statement. Circulation 136, 680–686.

Gardner, H., Kreidberg, J., Koteliansky, V., and Jaenisch, R. (1996). Deletion of integrin α1 by homologous recombination permits normal murine development but gives rise to a specific deficit in cell adhesion. Dev. Biol. 175, 301–313.

Gaubatz, S., Wood, J.G., and Livingston, D.M. (1998). Unusual proliferation arrest and transcriptional control properties of a newly discovered E2F family member, E2F-6. Proc. Natl. Acad. Sci. U. S. A. 95, 9190–9195.

González-Rosa, J.M., Sharpe, M., Field, D., Soonpaa, M.H., Field, L.J., Burns, C.E., and Burns, C.G. (2018). Myocardial Polyploidization Creates a Barrier to Heart Regeneration in Zebrafish. Dev. Cell 44, 433–446.e7.

Hashimoto, H., Yuasa, S., Tabata, H., Tohyama, S., Hayashiji, N., Hattori, F., Muraoka, N., Egashira, T., Okata, S., Yae, K., et al. (2014). Time-lapse imaging of cell cycle dynamics during development in living cardiomyocyte. J. Mol. Cell. Cardiol. 72, 241–249.

Hashimoto, H., Yuasa, S., Tabata, H., Tohyama, S., Seki, T., Egashira, T., Hayashiji, N., Hattori, F., Kusumoto, D., Kunitomi, A., et al. (2015). Analysis of cardiomyocyte movement in the developing murine heart. Biochem. Biophys. Res. Commun. 464, 1000–1007.

Hesse, M., Doengi, M., Becker, A., Kimura, K., Voeltz, N., Stein, V., and Fleischmann, B.K. (2018). Midbody Positioning and Distance Between Daughter Nuclei Enable Unequivocal Identification of Cardiomyocyte Cell Division in Mice. Circ Res 123, 1039–1052.

Jr, R.A., Wang, B.J., Quijada, P.J., Avitabile, D., Ho, T., Shaitrit, M., Chavarria, M., Firouzi, F., Ebeid, D., Monsanto, M.M., et al. (2018). Cardiomyocyte cell cycle dynamics and proliferation revealed through cardiac-specific transgenesis of fluorescent ubiquitinated cell cycle indicator (FUCCI). J. Mol. Cell. Cardiol. 127, 154–164.

Karbassi, E., Fenix, A., Marchiano, S., Muraoka, N., Nakamura, K., Yang, X., and Murry, C.E. (2020). Cardiomyocyte maturation: advances in knowledge and implications for regenerative medicine. Nat. Rev. Cardiol. 1–19.

Kimura, S.H., Ikawa, M., Ito, A., Okabe, M., and Nojima, H. (2001). Cyclin G1 is involved in G2/M arrest in response to DNA damage and in growth control after damage recovery. Oncogene 20, 3290–3300.

King, T., Bland, Y., Webb, S., Barton, S., and Brown, N.A. (2002). Expression of Peg1 (Mest) in the developing mouse heart: Involvement in trabeculation. Dev. Dyn. 225, 212–215.

Leone, M., Magadum, A., and Engel, F.B. (2015). Cardiomyocyte proliferation in cardiac development and regeneration: A guide to methodologies and interpretations. Am. J. Physiol. - Hear. Circ. Physiol. 309, H1237–H1250.

Liu, H., Zhang, C.-H., Ammanamanchi, N., Suresh, S., Lewarchik, C., Rao, K., Uys, G.M., Han, L., Abrial, M., Yimlamai, D., et al. (2019a). Control of cytokinesis by β-adrenergic receptors indicates an approach for regulating cardiomyocyte endowment. Sci. Transl. Med. 11, eaaw6419.

Liu, H., Zhang, C.H., Ammanamanchi, N., Suresh, S., Lewarchik, C., Rao, K., Uys, G.M., Han, L., Abrial, M., Yimlamai, D., et al. (2019b). Control of cytokinesis by β-adrenergic receptors indicates an approach for regulating cardiomyocyte endowment. Sci. Transl. Med. 11.

Liu, Z., Yue, S., Chen, X., Kubin, T., and Braun, T. (2010). Regulation of cardiomyocyte polyploidy and multinucleation by CyclinG1. Circ. Res. 106, 1498–1506.

Lun, A.T.L., McCarthy, D.J., and Marioni, J.C. (2016). A step-by-step workflow for low-level analysis of single-cell RNA-seq data [version 1; referees: 5 approved with reservations]. F1000Research 5.

Mahmoud, A.I., O’Meara, C.C., Gemberling, M., Zhao, L., Bryant, D.M., Zheng, R., Gannon, J.B., Cai, L., Choi, W.Y., Egnaczyk, G.F., et al. (2015). Nerves Regulate Cardiomyocyte Proliferation and Heart Regeneration. Dev. Cell 34, 387–399.

Mollova, M., Bersell, K., Walsh, S., Savla, J., Das, L.T., Park, S., Silberstein, L.E., Dos Remedios, C.G., Graham, D., Colan, S., et al. (2013). Cardiomyocyte proliferation contributes to heart growth in young humans. Proc. Natl. Acad. Sci. U. S. A. 110, 1446–1451.

Moon, Y.S., Park, S.K., Kim, H.T., Lee, T.S., Kim, J.H., and Choi, Y.S. (2010). Imprinting and expression status of isoforms 1 and 2 of PEG1/MEST gene in uterine leiomyoma. Gynecol. Obstet. Invest. 70, 120–125.

Pedersen, I.S., Dervan, P.A., Broderick, D., Harrison, M., Miller, N., Delany, E., O’Shea, D., Costello, P., McGoldrick, A., Keating, G., et al. (1999). Frequent loss of imprinting of PEG1/MEST in invasive breast cancer. Cancer Res. 59, 5449–5451.

Picelli, S., Björklund, Å.K., Faridani, O.R., Sagasser, S., Winberg, G., and Sandberg, R. (2013). Smart-seq2 for sensitive full-length transcriptome profiling in single cells. Nat. Methods 10, 1096–1100.

Porrello, E.R., Mahmoud, A.I., Simpson, E., Hill, J.A., Richardson, J.A., Olson, E.N., and Sadek, H.A. (2011). Transient regenerative potential of the neonatal mouse heart. Science (80-.). 331, 1078–1080.

Pozo, P., and Cook, J. (2016). Regulation and Function of Cdt1; A Key Factor in Cell Proliferation and Genome Stability. Genes (Basel). 8, 2.

Puente, B.N., Kimura, W., Muralidhar, S.A., Moon, J., Amatruda, J.F., Phelps, K.L., Grinsfelder, D., Rothermel, B.A., Chen, R., Garcia, J.A., et al. (2014). The Oxygen-Rich Postnatal Environment Induces Cardiomyocyte Cell-Cycle Arrest through DNA Damage Response. Cell 157, 565–579.

Quaife-Ryan, G.A., Sim, C.B., Ziemann, M., Kaspi, A., Rafehi, H., Ramialison, M., El-Osta, A., Hudson, J.E., and Porrello, E.R. (2017). Multicellular transcriptional analysis of mammalian heart regeneration. Circulation 136, 1123–1139.

Ramsköld, D., Wang, E.T., Burge, C.B., and Sandberg, R. (2009). An abundance of ubiquitously expressed genes revealed by tissue transcriptome sequence data. PLoS Comput. Biol. 5, 1–11.

Raulf, A., Horder, H., Tarnawski, L., Geisen, C., Ottersbach, A., Röll, W., Jovinge, S., Fleischmann, B.K., and Hesse, M. (2015). Transgenic systems for unequivocal identification of cardiac myocyte nuclei and analysis of cardiomyocyte cell cycle status. Basic Res. Cardiol. 110, 489.

Sakaue-Sawano, A., Kurokawa, H., Morimura, T., Hanyu, A., Hama, H., Osawa, H., Kashiwagi, S., Fukami, K., Miyata, T., Miyoshi, H., et al. (2008). Visualizing spatiotemporal dynamics of multicellular cell-cycle progression. Cell 132, 487–498.

Senyo, S.E., Steinhauser, M.L., Pizzimenti, C.L., Yang, V.K., Cai, L., Wang, M., Wu, T.-D., Guerquin-Kern, J.-L., Lechene, C.P., and Lee, R.T. (2013). Mammalian heart renewal by pre-existing cardiomyocytes. Nature 493, 2–6.

Stopp, S., Grundl, M., Fackler, M., Malkmus, J., Leone, M., Naumann, R., Frantz, S., Wolf, E., Eyss, B. von, Engel, F.B., et al. (2017). Deletion of Gas2l3 in mice leads to specific defects in cardiomyocyte cytokinesis during development. Proc Natl Acad Sci U S A 114, 8029–8034.

Storvall, H., Ramsköld, D., and Sandberg, R. (2013). Efficient and Comprehensive Representation of Uniqueness for Next-Generation Sequencing by Minimum Unique Length Analyses. PLoS One 8.

Uva, G.D., Aharonov, A., Lauriola, M., Kain, D., Yahalom-ronen, Y., Carvalho, S., Weisinger, K., Bassat, E., Rajchman, D., Yifa, O., et al. (2015). ERBB2 triggers mammalian heart regeneration by promoting cardiomyocyte dedifferentiation and proliferation.

Wei, K., Serpooshan, V., Hurtado, C., Diez-Cuñado, M., Zhao, M., Maruyama, S., Zhu, W., Fajardo, G., Noseda, M., Nakamura, K., et al. (2015). Epicardial FSTL1 reconstitution regenerates the adult mammalian heart. Nature.

Windmueller, R., Leach, J.P., Babu, A., Zhou, S., Morley, M.P., Wakabayashi, A., Petrenko, N.B., Viatour, P., and Morrisey, E.E. (2020). Direct Comparison of Mononucleated and Binucleated Cardiomyocytes Reveals Molecular Mechanisms Underlying Distinct Proliferative Competencies. Cell Rep. 30, 3105–3116.e4.

Xin, M., Kim, Y., Sutherland, L.B., Murakami, M., Qi, X., McAnally, J., Porrello, E.R., Mahmoud, a. I., Tan, W., Shelton, J.M., et al. (2013). Hippo pathway effector Yap promotes cardiac regeneration. Proc. Natl. Acad. Sci. 110, 13839–13844.

Young, M.D., Wakefield, M.J., Smyth, G.K., and Oshlack, A. (2010). Gene ontology analysis for RNA-seq: accounting for selection bias. Genome Biol. 11.

Yue, F., Cheng, Y., Breschi, A., Vierstra, J., Wu, W., Ryba, T., Sandstrom, R., Ma, Z., Davis, C., Pope, B.D., et al. (2014). A comparative encyclopedia of DNA elements in the mouse genome. Nature 515, 355–364.

