## Supplementary figures and images for "Identification and characterization of distinct cell cycle stages in cardiomyocytes using the FUCCI transgenic system"

### Supplementary Figure 1

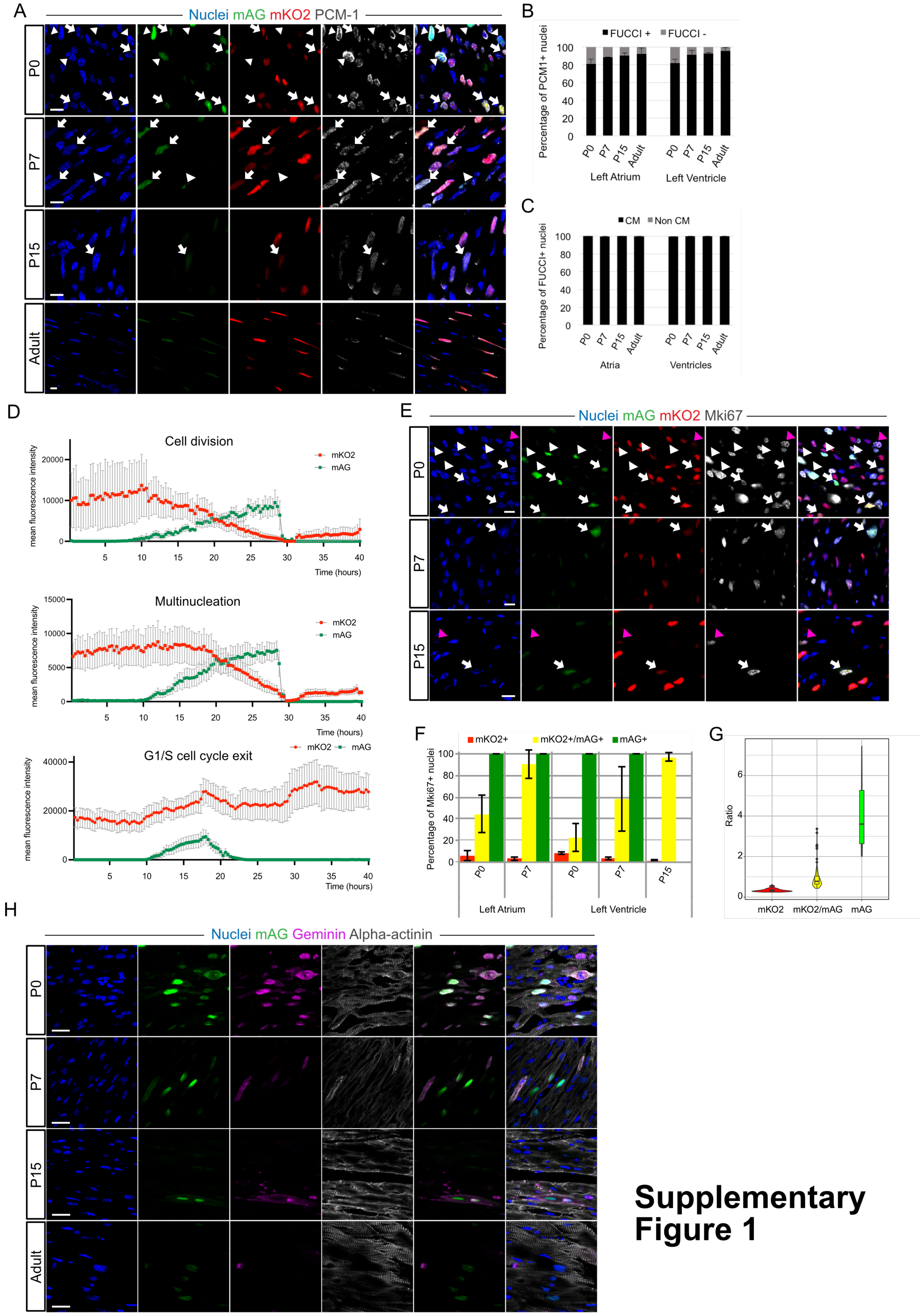


#### Supplementary Figure 1

### Supplementary Figure 2

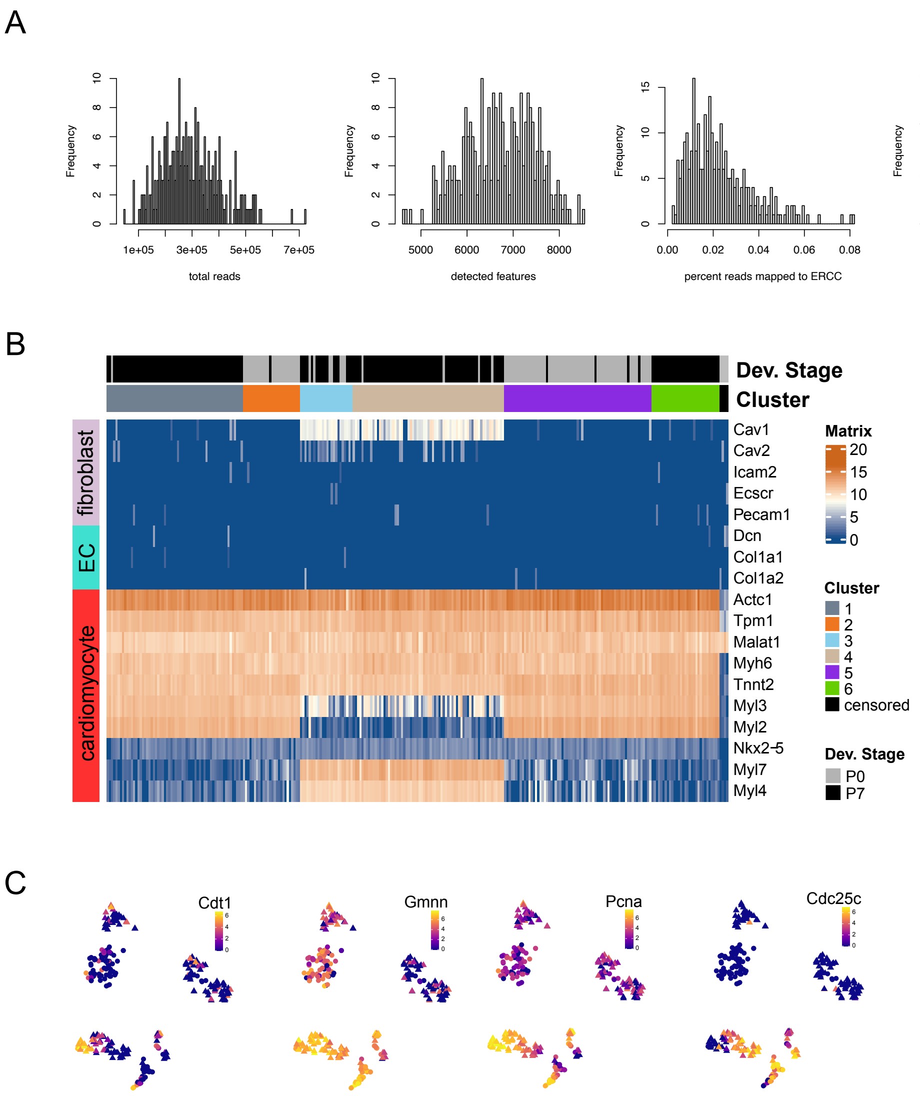


Supplementary Figure 2

### Supplementary Figure 3

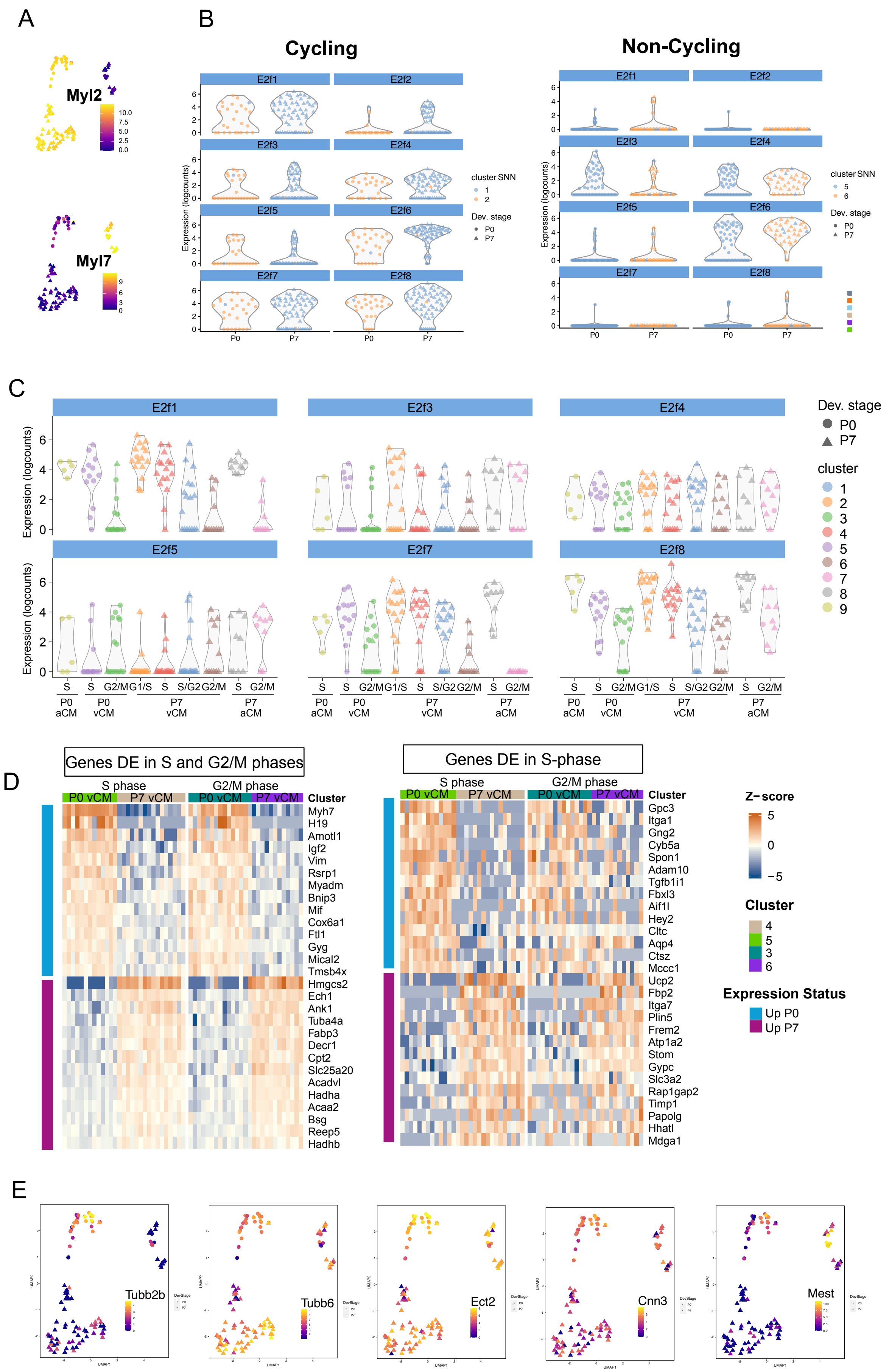


**Supplementary Figure 3**
